# Tardigrades’ cytoplasmic abundant heat soluble proteins serve as membrane protectors during dehydration

**DOI:** 10.1101/2025.04.14.648834

**Authors:** Claire K. Zhang

## Abstract

Tardigrades are microscopic organisms with extraordinary tolerance to environmental stresses such as desiccation and thus offer unique solutions to bio-preservation, anti-aging, and interstellar travel. Recent studies revealed a collection of cytoplasmic and secretory abundant soluble proteins (CAHSs and SAHSs respectively) contributing to tardigrades’ extreme resilience. Using computational tools, I examined 39 CAHSs and 28 SAHSs from three representative tardigrade species. Both protein families possess a conserved central region and two highly variable terminal regions. Phylogenetic analysis suggests that CAHSs and SAHSs have distinct sequences despite functional similarity. AlphaFold predicts that CAHSs’ central region forms a long and amphiphilic α-helix whereas SAHSs’ folds into β-barrel. Since dehydration leads to the increase of intracellular protein concentration, I used AlphaFold to simulate CAHS oligomerization and find that they preferably dimerize via their central helix motifs. Examination of CAHS dimers reveals a strong inter-helix interaction. The dimerized anti-parallel helix bundle has hydrophobic and hydrophilic sides, resembling lipid-interacting proteins like ApoE. Empirical tests using mammalian fibroblast cells expressing the representative RvCAHS3 show that CAHSs concentrated on intracellular membranes instead of in proteinaceous condensates during dehydration. Moreover, RvCAHS3 significantly improves cell survival as measured by the stimulation-evoked release of Ca^2+^ from membrane-enclosed internal stores like endoplasmic reticulum. Taken together, these results suggest that CAHS inclines to dimerize and further forms a mesh on intracellular membranes to reinforce them during environmental stresses. By so doing, CAHSs can protect the integrity and the functionality of membrane-enclosed organelles. This finding implicates new strategies to preserve biomolecules, cells, and tissues under challenging conditions or for easy transportation.

## Introduction

Tardigrades, commonly known as water bears or moss piglets, are microscopic invertebrates comprising the distinct phylum Tardigrada (1). They have a segmented body with four pairs of legs and live in marine or freshwater environments as well as semi-terrestrial habitats (1). Tardigrades are renowned for withstanding extreme environments including desiccation, extreme temperatures, and cosmic radiation (2). Most notably, they can remain lifeless in a vacuum for decades until rehydration (3). Thus, tardigrades are exceptional models for studying anhydrobiosis (i.e., an organism loses almost all its water and enters a state of reversible ametabolism) (4). For most organisms, dehydration causes hyperosmosis and damages cellular structures, leading to cell deformation and eventually death. Tardigrades, on the other hand, have various protective measures for such challenges.

Recent studies have found that two protein families, cytosolic and secretory abundant heat soluble proteins (CAHSs and SAHSs), serve as intracellular and extracellular protectants in tardigrades, respectively (5). Early research had concluded that both were intrinsically disordered and vitrified upon dehydration, hypothetically sequestering biomolecules, organelles, and cellular apparatuses (5). Lately, computational as well as empirical studies have indicated that those proteins have a defined tertiary structure, at least partially (6-9). Moreover, the latest investigations have shown that CAHSs are protective for protein complexes and organelles (6) whereas SAHSs shelter extracellular biomolecules and structures (10). To better appraise their protective mechanisms, especially in case of desiccation, I have employed *in silico* and *in vitro* tests with a focus on the oligomerization of CAHSs owing to dehydration. As more and more tardigrades have undergone genomic sequencing (11, 12), more and more CAHS and SAHS gene sequences have been deposited in GenBank and become available to the general public. More importantly, sequence-based structure modeling fueled by the latest development in artificial intelligence has become highly reliable (e.g., AlphaFold 3.0 achieved over 97% accuracy in predicting protein complex) (13). In conjunction with computer simulation, empirical tests in model systems like cultured cells yield insights bridging proteins’ functionality with their structure.

## Materials and Methods

All nucleotide and protein sequences used for this study were obtained from GenBank and UniProt using keyword searching (i.e., abundant heat soluble protein, CAHS, or SAHS) and filtered by selected tardigrade species. All protein sequence alignments were performed using constraint-based multiple alignment tools (i.e., COBALT) (14) available from the National Center for Biotechnology Information. The default alignment parameters were used. To generate phylogenetic trees based on the sequence alignments, I used the ETE3 toolkit with default settings available from GenomeNet (www.genome.jp). All structural models of CAHSs and SAHSs were generated using AlphaFold 3.0 (13). The default settings were used to ensure a fairness to all proteins. Resulting structures were downloaded and visualized using UCSF Chimera program (15).

All chemical reagents were acquired from Thermo Fisher Scientific unless specified. 3T3 cells were gifted from Dr. Henriette van Praag. All DNA plasmids were acquired from Addgene. DNA extraction and purification were completed using MaxiPrep kit from Zymo Research. DNA transfection to 3T3 cells were done using Lipofectamine. Confocal fluorescence imaging was carried out using Nikon A1R confocal system, and Ca^2+^-imaging were conducted with a Nikon Ti-E microscope controlled by μManager (16). Image analyses were executed using FIJI (17). Statistical analyses and plots were done using Excel and/or Prism.

## Results

From among ∼1,500 species of tardigrades, I selected three representative ones: *Ramazzottius varieornatus* (*Rv*) is best known for its extremotolerance (18); *Hypsibius exemplaries* (*He*) is the most studied for evolutionary biology and astrobiology (19); and *Paramacrobiotus metropolitanus* (*Pm*) is a popular genetic model (20). Most of all, their genomes have been sequenced and almost all extremotolerance-related genes have been identified and deposited in public-accessible databases like GenBank (21). After exhaustive searching, I acquired all DNA and protein sequences of CAHSs and SAHSs in those three species (Tables S1 and S2).

First, the protein sequences of 39 CAHSs and 28 SAHSs were aligned using COBALT with default settings. A distinct consensus region was observed in each group (Figure 1). In CAHSs, the conserved region of approximately 130 amino acid residues was flanked by N- and C-terminal regions with highly variable lengths and amino acid compositions (Figure 1A). Notably, the most conserved amino acid residues (shown in red in Figure 1A) are hydrophilic (i.e., charged or polar) and distributed evenly across the consensus region (Figure S1A). In SAHSs, the conserved region is made of approximately 100 amino acid residues and is closer to the C-terminals (Figure 1B). Different from that of CAHSs, it is less consistent and can be segmented into three subregions of shorter and variable sequences, about 20 ∼ 40-residues long. The most conserved amino acid residues in the conserved regions of SAHSs are either hydrophilic or hydrophobic (Figure S1B). The phylogenetic tree plots suggest that CAHSs are more conserved across different species because there are more species than gene differences between the neighboring CAHSs (Figure S2A). Notably, PmCAHS89226-like was found to be phylogenetically distant from all other 38 CAHS, consistent with its sequence alignment, indicating an incorrect categorization. As for SAHSs, the closest genes are always from the same species (Figure S2B). Even combined together, CAHSs and SAHSs form two separate branches (Figure S2C). In summary, the sequence analyses suggests that CAHSs and SAHSs are two very different families of proteins despite their shared names and functional similarity. Furthermore, their conserved regions have very different amino acid compositions, implicating differences in protein structure and function.

**Figure 1.**
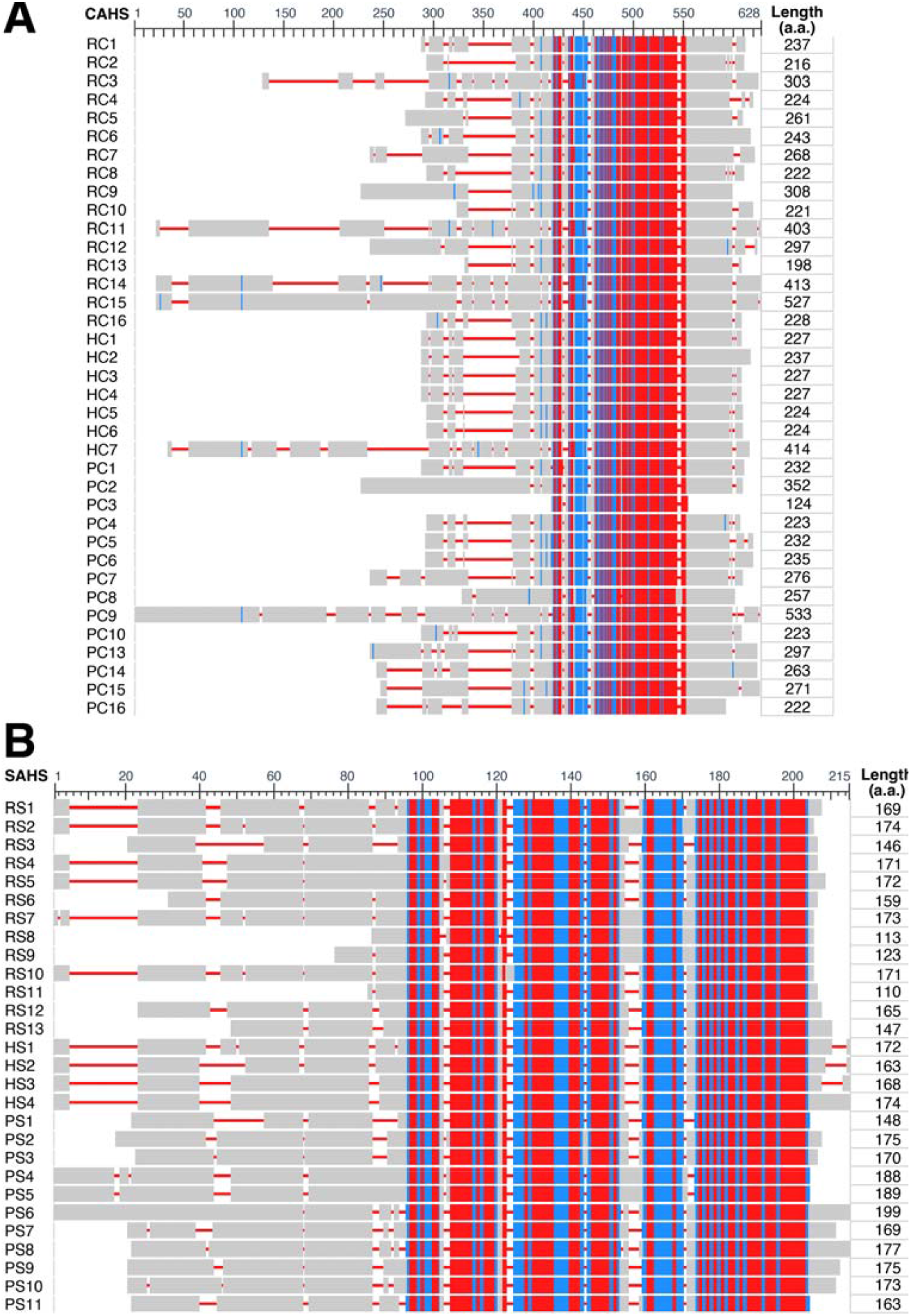
Protein sequence alignment identifies highly conserved central motifs and highly variable terminal regions in CAHSs and SAHSs. Alignments of CAHSs (**A**) and SAHSs (**B**) were generated by COBALT. Red represents highly conserved amino acid residues, blue for less conserved ones, and gray for highly variable ones. The conserved central motif for CAHSs is about 120 amino acids long, and SAHSs’ is about 100 amino acids long. RC, Rv CAHS; HC, He CAHS; PC, Pm CASH. They are all numbered in the same order as that of Figure S1 and Table S1 & S2.

To model CAHSs and SAHSs, the AI-based AlphaFold was used because its latest version (3.0) offers unprecedented accuracy and reliability, especially in predicting protein complex including oligomers (13). Consistent with the sequence alignment, all of CAHSs’ long conserved regions formed a single α-helix (Figure 2A and S3A) except for PmCAHS89226-like, again suggesting that it was miscategorized. So, it is excluded from all analyses thereafter. In case of SAHSs, their conserved regions form several consecutive β-sheets with α-helixes or coils in between (Figure S3B). Unlike CAHSs, there is no outlier in SAHSs, again consistent with the phylogenetic analysis result. As for the highly variable N- and C-terminal regions in CAHSs and SAHSs, they are generally deemed to be disordered by AlphaFold with low predicted local distance difference test score (plDDT) (Figure 2A and S3).

**Figure 2.**
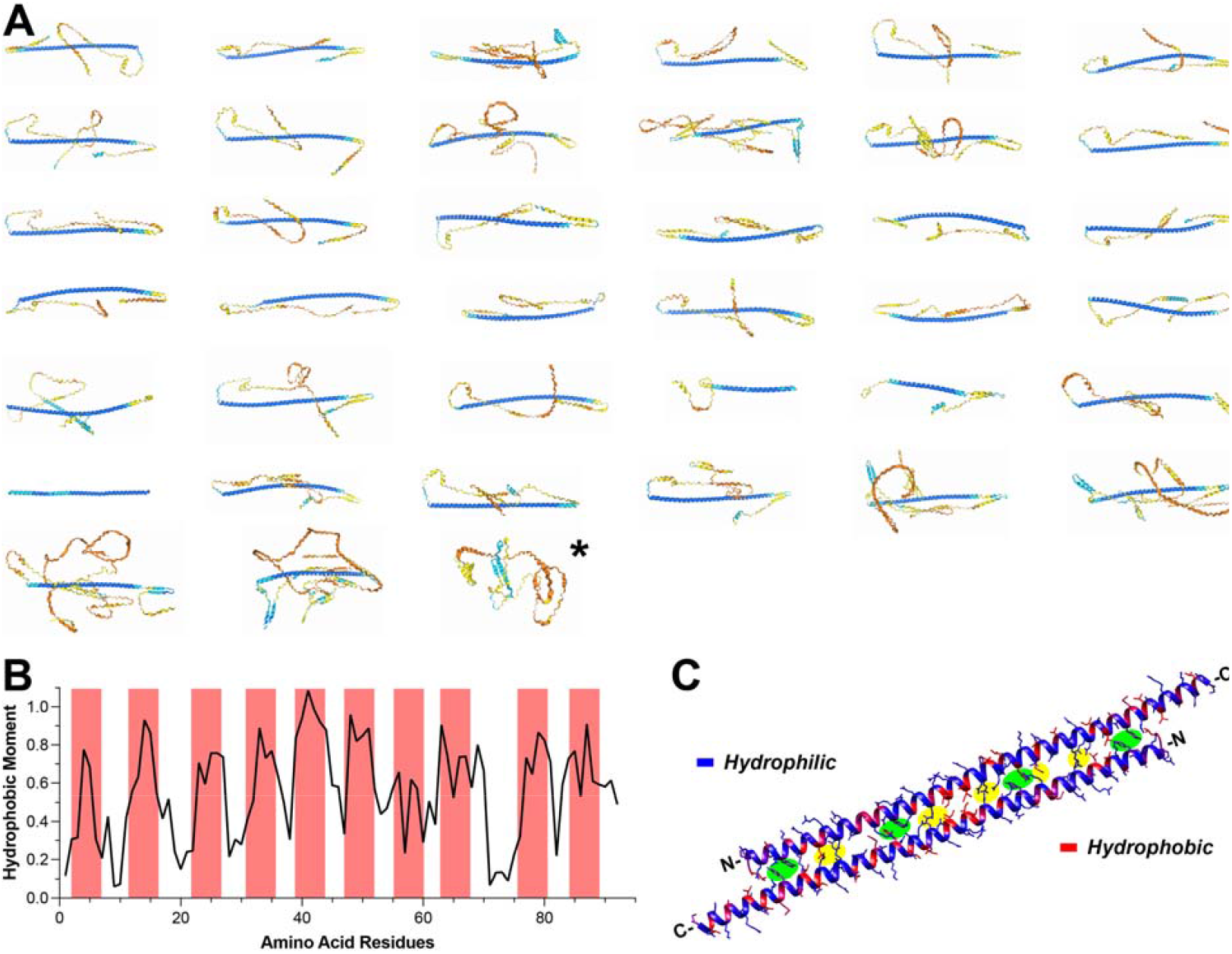
CAHS’s central motif forms long helix. **A**, the 3D structures of CAHS monomers predicted by AlphaFold. Blue indicates high confidence for the highly ordered central helix whereas yellow/orange indicates low confidence for the disorded terminals. (*, PmCAHS89226-like) **B**, the hydrophobic moment plot of RvCAHS3’s central motif shows hydrophobic residues are arranged in a periodic manner. **C**, two central motifs of RvCAHS3 form an anti-parallel dimer via electrostatic interactions (highlighted in green) and π-π stacking (highlighted in yellow). Most hydrophobic residues face outside.

Using RvCAHS3 as an example, the hydrophobic moments along the conserved central regions were calculated, which clearly exhibits periodic peaks (Figure 2B). This is consistent with the evenly distributed hydrophobic amino acid residues shown in Figure S1A and predicts that the α-helix is likely amphiphilic. Dehydration effectively concentrates biomolecules and thus promotes CAHS/SAHS oligomerization inside and outside of cells. So, AlphaFold 3.0 was employed to model the formation of CAHS and SAHS oligomers. For example, when a second helix of RvCAHS3 conserved regions was introduced, AlphaFold yielded a dimer of two helixes in an anti-parallel fashion (Figure 2C). A close examination of inter-peptide interactions revealed multiple electrostatic interactions and π-π stacking along the interface of the two helixes (highlighted in Figure 2C), indicating a high stability of such dimer. Moreover, the hydrophobic (red) and hydrophilic (blue) segments of both helixes were well aligned in the dimer (Figure 2C), very much reminiscent to helix bundles in the lipid-binding proteins like Apolipoprotein E (9, 22).

Like RvCAHS3 (Figure 3A1), all CAHS dimers and trimers were formed in an anti-parallel fashion with moderate increase or decrease of prediction confidence (i.e., predicted template modeling score, pTM) in comparison to monomers (Figure 3A2). For the central helix motif alone, the overall confidence scores were the highest for dimers but dropped sharply for trimers (Figure 3B). Due to the low prediction scores of the disordered regions obscuring interactions between the central helix motifs, the consensus regions of CAHSs were used for the subsequent modeling of CAHS oligomerization thereafter. Figure 3C illustrates a clear trend of decrease in pTM as oligomerization progresses. Due to the strong binding in the dimers and the high confidence in dimer prediction, it is very likely that desiccation promotes CAHSs to dimerize and the dimers connect to each other to form a protective mesh on lipid membranes. I further speculate that the presence of such a CAHS oligomer cover can prevent merging or collapsing of membranes to each other and consequently prevent the breakdown of membrane-enclosed organelles, a requisite for cell survival.

**Figure 3.**
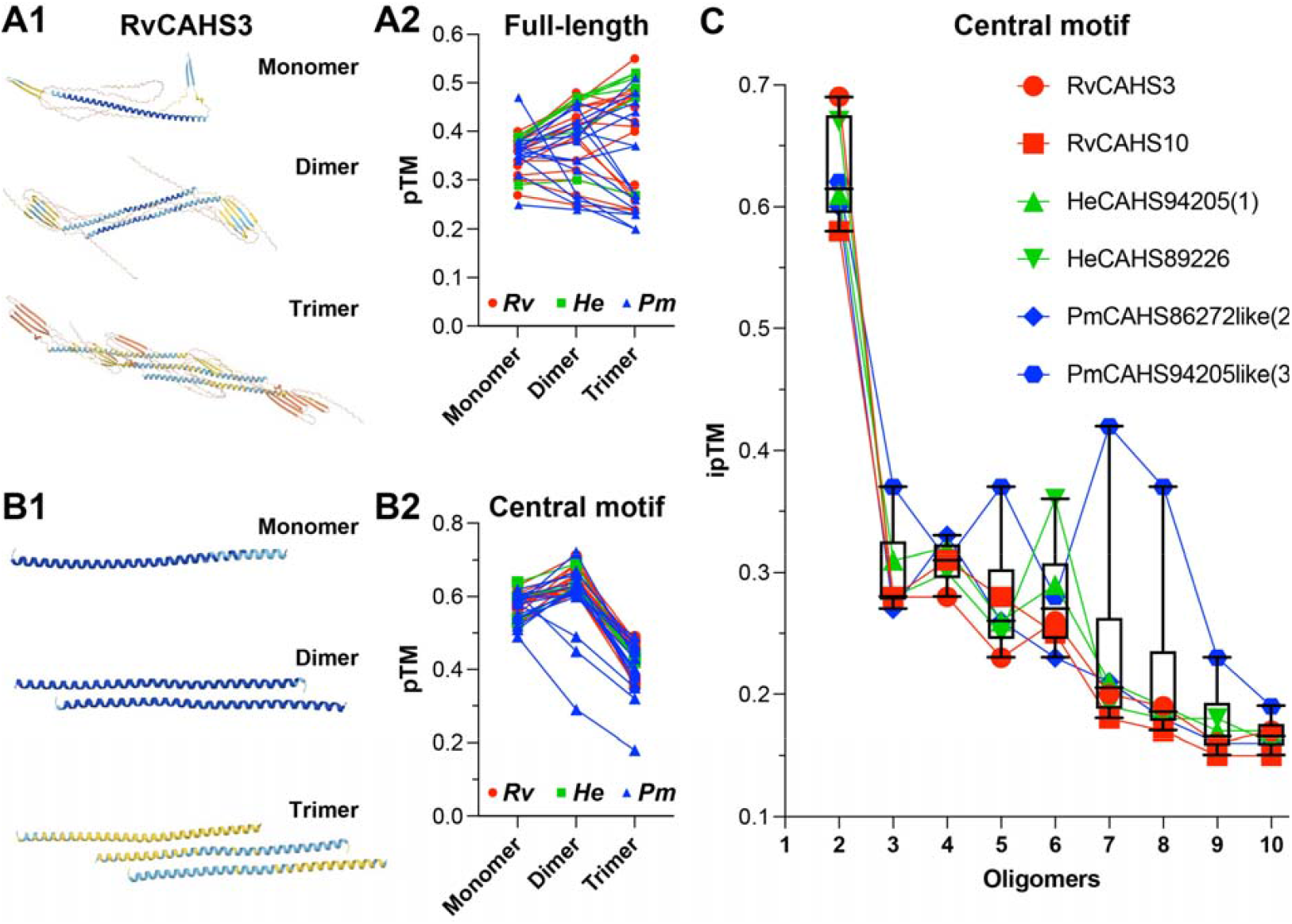
A helix dimer is the most likely form during CAHS oligomerization. **A**, predicted structures of full-length RvCAHS3 in mono-, di-, and trimer forms (**A1**) and the prediction confidence (measured as pTM) for the mono-, di-, and trimers of all 39 full-length CAHSs (**A2**), which lacks an overall trend of changes. **B**, predicted structures of RvCAHS3 central motif in mono-, di-, and trimer forms (**B1**) and the prediction confidence (measured as pTM scores) of the central motifs of all 39 CAHSs (**B2**), which shows an overall increase for dimers and a drastic decrease for timers. **C**, the box and whisker plot combined with point-line plot shows the trend of prediction confidence (measured as ipTM) during the oligomerization (from dimers to decamers) of the central motifs of six representative CAHSs. There is a progressive decrease from dimers to decamers.

In order to empirically test that idea, 3T3 cells (i.e., immortalized mouse embryonic fibroblast cells) growing on Matrigel-coated glass coverslips were transfected with a mammalian-expressing plasmid encoding RvCAHS3, which is tagged with green fluorescent protein (i.e., CAHS3-AcGFP1) for detection by fluorescence microscopes (23). About 1 day after the transfection, more than 80% cells expressed CAHS3-AcGFP1 (estimated by AcGFP1 fluorescence). In order to simulate dehydration, cells growing on the coverslips were air-dried in a laminar flow cabinet at room temperature (∼25°C) for different periods of time (i.e., 0, 1, 2, 5, 10, and 20 minutes). Previous studies suggested that CAHSs underwent gel-transition or liquid-liquid phase separation (LLPS) upon dehydration-like treatments (9, 23). To test if CAHS3 does that in 3T3 cells, they were co-transfected with a DsRed-expressing plasmid. It is well documented that DsRed inclines to aggregate, vitrify, and form LLPS-like protein condensates (24). To visualize membrane-enclosed organelles, those transfected cells were incubated with FM4-64, a far-red fluorescent dye that can reversibly insert into lipid bilayers and label intracellular membranes after being endocytosed. After loading, the FM4-64 remaining on the cell surface membrane was readily washed off by a 5-minute perfusion with dye-free normal Tyrode’s solution (in mM: NaCl, 140; KCl, 2; CaCl_2_, 2; MgCl_2_, 2; HEPES, 10; D-Glucose, 10mM. pH7.35; 305 Osm/L). Figure 4A and supplementary movies exemplify such triple-labeled 3T3 cells (blue represents cell membrane; green is CASH3; and red indicates protein condensates). After 5-min air drying in a laminar flow cabinet, the majority of AcGFP1 fluorescence was found to be colocalized with that of FM4-64 but not DsRed (Figure 4A). Consistent with the observation, there is a statistically significant correlation between AcGFP1 and FM4-64 signals but not those of DsRed (Figure 4B&C), meaning the larger membrane areas were the more CAHS3 associated with them. This result suggests that dehydration drove most CAHS3 onto intracellular membranes instead of LLPS-related protein condensates.

**Figure 4.**
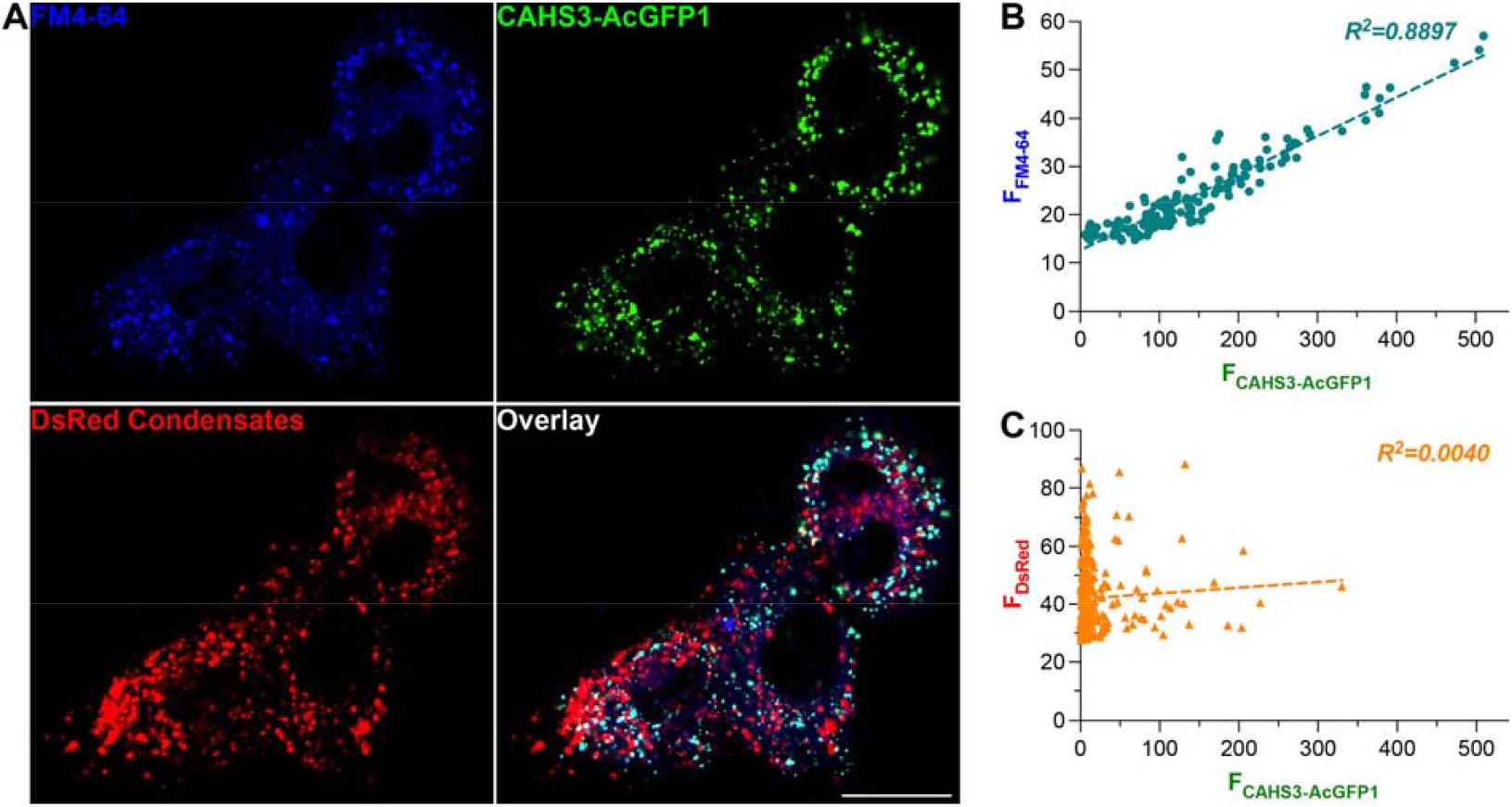
During dehydration, CAHS is mostly associated with cell membranes rather than proteinaceous condensates. **A**, sample confocal fluorescence images of DsRed and CAHS3-AcGFP1-expressing cells pre-loaded with FM4-64 and air-dried at 25°C for 5 minutes. Scale bar, 20μm. **B**, a scatter plot of FM4-64 and CAHS3-AcGFP1 fluorescence intensities within the puncta defined by FM4-64 (i.e., membrane-bound organelles). The two values are significantly correlated (*p* < 0.0001), i.e., CAHS3-AcGFP1 colocalized with the membranes during dehydration. **C**, a scatter plot of DsRed and CAHS3-AcGFP1 fluorescence intensities within the puncta defined by DsRed (i.e., proteinaceous condensates). The two values are not correlated (*p* = 0.3114), i.e., CAHS3-AcGFP1 did not co-condense with DsRed during dehydration.

Next, Ca^2+^-imaging was used to test if CAHS3 protects membrane-enclosed organelles and made cells more resilient to dehydration. For that, transfected 3T3 cells and the sham controls were pre-loaded with a cell membrane-permeable red fluorescent Ca^2+^-indicator (i.e., X-Rhod-1AM) (25) before they were air dried. Immediately after drying, those cells were rehydrated and continuously perfused with the normal Tyrode’s solution. During imaging, 50 μM ATP was used to stimulate those stressed cells. Such ATP stimulation usually causes the release of Ca^2+^ from internal stores like endoplasmic reticulum (ER, a major membrane-enclosed organelles), which tests not only cell responsiveness (i.e., viability) but also the integrity of membrane-enclosed organelles. Figure 5A shows that longer air-drying caused less cells to respond in both CAHS3 group and the control. However, the CAHS3 expression made more cells responding to the ATP stimulation than the sham control (Figure 5B). Statistical significance was reached at 2, 5, 10, and 20 minutes. More importantly, the average amplitude of such Ca^2+^ response was much higher in the CAHS3-expressing group than the control (Figure 5C), supporting the idea that the internal Ca^2+^ stores in CAHS3-expressing cells were more robust than those in the controls. Taken together, the membrane association of CAHS3 and the better maintained organelles all suggest that CAHS3 reinforces intracellular membranes and effectively enhances mammalian cell survival during prolonged dehydration.

**Figure 5.**
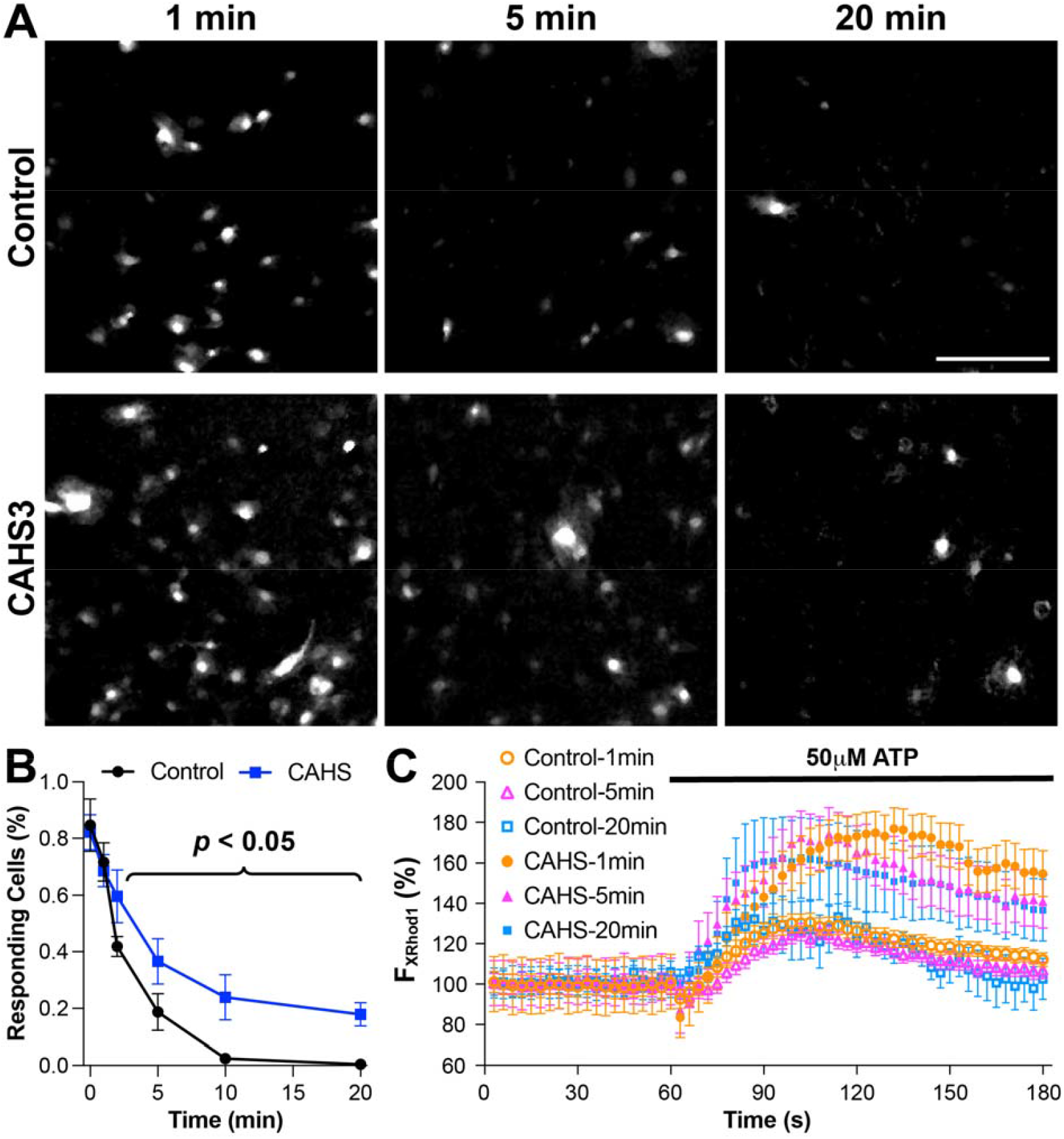
CAHS3 increased cell survival during dehydration. **A**, sample fluorescence images of control and CAHS3-expressing 3T3 cells still responding to 50μM ATP stimulation (visualized by X-Rhod-1AM, a red Ca^2+^ indicator) after 1, 5, and 20-min air-drying at 25°C. Scale bar, 100μm. **B**, percentage of responding cells (mean ± SEM) after air-drying at 25°C. The CAHS3-expressing group had more responding cells than the control group starting from the 5-min time point (all *p* <0.05). **C**, relative change of X-Rhod-1 fluorescence in individual responding cells shows significant increase after the application of ATP. The average fluorescence intensity in every responding cell at every time point was normalized to its initial value in the first image.

## Discussion

The fascinating ability of tardigrades to sustain and survive extreme environments such as the vacuum of space ignites great interest in using them as model organisms to study biological mechanisms for bio-preservation, anti-aging, and space travel (1-3). The physiological basis for tardigrades’ extremotolerance is anhydrobiosis, for which intrinsically disordered proteins, namely CAHSs and SAHSs, are known to be essential. As more and more CAHSs and SAHSs have been discovered in different species of tardigrades (18-21), it becomes clear that both of them are indispensable for the preservation of intracellular and extracellular structures and functions during anhydrobiosis (5, 6, 10). Previously, these unstructured proteins were believed to work as absorbents for intra- and extracellular biomolecules (5). However, it is puzzling how a single mechanism can deal with very different needs by intracellular and extracellular apparatuses. Furthermore, significant differences in protein sequences and subcellular localizations between CAHSs and SAHSs speak against the notion of a shared mechanism between the two.

To better understand CAHSs and SAHSs, I started with structural analysis. As proteins’ functions are largely determined by their peptide sequences, I collected all CAHSs and SAHSs sequences in three representative tardigrade species from GenBank. The sequence alignments unveiled highly conserved regions in both CAHSs and SAHSs, which are significantly different from each other (Figure 1), which is confirmed by their separation in the phylogenetic trees (Figure S2C). Next, AlphaFold 3.0 consistently predicted a single α-helix for the consensus region of CAHSs (Figures 2, 3, and S3A&C) and a mix of β-sheets and short α-helixes for that of SAHSs (Figure S3B&D), which suggests that neither CAHSs nor SAHSs are completely disordered. Given their structural difference, CAHSs and SAHSs very likely act differently for cell protection.

Due to their unique and highly stable helical dimers (Figure 3), I focused on CAHSs. Intriguingly, the highly conserved hydrophilic amino acid residues and repeated hydrophobic moments (Figure 2B&C) result in the periodic hydrophobicity and hydrophilicity across the helical bundle, a characteristic structure found in lipid-binding proteins like ApoE (9, 22). This indicates that CAHS dimers favorably interact with lipid membranes. Furthermore, the connection of CAHS dimers via their unstructured terminal regions likely promotes the formation of CAHS-dimer networks covering cell membranes. This prediction is supported by the experimental observation that RvCAHS3 mostly co-localizes with intracellular membrane label (i.e., FM4-64) upon dehydration (Figure 4). Although it differs from a previous observation that CAHSs vitrified upon environmental challenge (9), my result aligns with the report that CAHSs inclined to form network of oligomers during hyperosmotic stress (23). Hence, I propose that dehydration promotes the formation of a web of CAHS dimers, which attaches to intracellular membranes and creates a barrier to prevent the collapsing or merging of intracellular membranes. By doing so, CAHSs can help membrane-enclosed organelles to retain their integrity when cells undergo desiccation. Again, this idea is supported by the observation that CAHS3-expressing 3T3 cells exhibited significantly better Ca^2+^ response than the control after prolonged dehydration (>2 minutes) (Figure 5).

Due to the constraints of AlphaFold, the oligomer modeling could not account for changes in biomolecule mixing, ion concentration, or other extracellular and intracellular changes during dehydration. Additionally, there were unaccounted errors due to the limitations in AI algorithms and training datasets used by AlphaFold. Nevertheless, the fact that AlphaFold consistently predicts the helical central motifs and helix dimers for most CAHSs reassures the structural prediction. The cell-based assays so far only investigated RvCAHS3 in the cytoplasm. Thus, it is worthwhile to expand such empirical study to other CAHSs from different species of tardigrades or bearing structural difference from RvCAHS3. It is also interesting to investigate if such a mechanism by RvCAHS3 can protect cell surface membrane, which can be achieved by adding a secretory signaling sequence to RvCAHS3, relocating it to extracellular spaces. In addition, alternative challenges like hyperosmotic stress or different types of cells like the more fragile neurons can be used to explore the protection capacity of CAHSs and SAHSs. Future research on their protective mechanisms should be extended to whole animals using model organisms such as *C. elegans*, which is certainly more informative for translational applications. Last but not least, such hybrid studies combing computational and empirical analyses can be applied to intrinsically disordered proteins native to mammalian cells (e.g., late embryogenesis abundant proteins) to investigate and improve their protective effects for clinical use.

## Supporting information

Calcium-imaging of control 3T3 cells after 1-min dehydration

Calcium-imaging of control 3T3 cells after 5-min dehydration

Calcium-imaging of control 3T3 cells after 20-min dehydration

Calcium-imaging of CAHS-expressing 3T3 cells after 1-min dehydration

Calcium-imaging of CAHS-expressing 3T3 cells after 5-min dehydration

Calcium-imaging of CAHS-expressing 3T3 cells after 20-min dehydration

Example 1 of 3D-reconstructured live-cell confocal images of 3T3 cells expressing DsRed and RvCAHS-AcGFP1 and preloaded with FM4-64

Example 2 of 3D-reconstructured live-cell confocal images of 3T3 cells expressing DsRed and RvCAHS-AcGFP1 and preloaded with FM4-64

Example 3 of 3D-reconstructured live-cell confocal images of 3T3 cells expressing DsRed and RvCAHS-AcGFP1 and preloaded with FM4-64

## Acknowledgment

I would like to thank Dr. Cristina Fenollar Ferrer in the Stiles-Nicholson Brain Institute at Florida Atlantic University for mentoring me and supporting my research.

## Supporting information

**S1 Fig. Characterization of amino acid residues within the consensus sequences of CAHSs (A) and SAHSs** (**B)**. Red, hydrophobic; blue, hydrophilic; gray, neutral.

**S2 Fig. Phylogenetic trees of CAHSs (A), SAHSs (B), and both (C), generated by ETE3**.

**S3 Fig. Structural prediction of 39 CAHSs and 28 SAHSs in monomer (A and B)**. The prediction confidence is measured as the predicted local distance difference test (plDDT). Dark blue indicates high confidence (i.e., plDDT > 90), blue to cyan indicates good confidence (i.e., 90 > plDDT > 70), yellow indicates low confidence (i.e., 70 > pTM > 50), and orange indicates very low confidence (i.e., pTM < 50).

**S1 Table. All CAHSs used in this study**.

**S2 Table. All SAHSs used in this study**.

**S1 Files. Ca**^**2+**^**-imaging of Controls after 1, 5, and 20-minute airdry (3 AVI files)**.

**S2 Files. Ca**^**2+**^**-imaging of CAHS3-expressing cultures after 1, 5, and 20-minute airdry (3 AVI files). S3 Files. 3D view of reconstructed confocal images showing CAHS3-AcGFP1 (green), DsRed (red), and FM4-64 (blue) in 3T3 cells after 5-minute airdry (3 AVI files)**.

