## Supplementary figures and tables for "Tardigrades’ cytoplasmic abundant heat soluble proteins serve as membrane protectors during dehydration"

Full title:

Short title:

CAHS protects cell membranes during dehydration

Claire K. Zhang[1]*

[1]Brentwood High School, Brentwood, Tennessee, United States of America

* Corresponding author

### I. Supplementary Figures

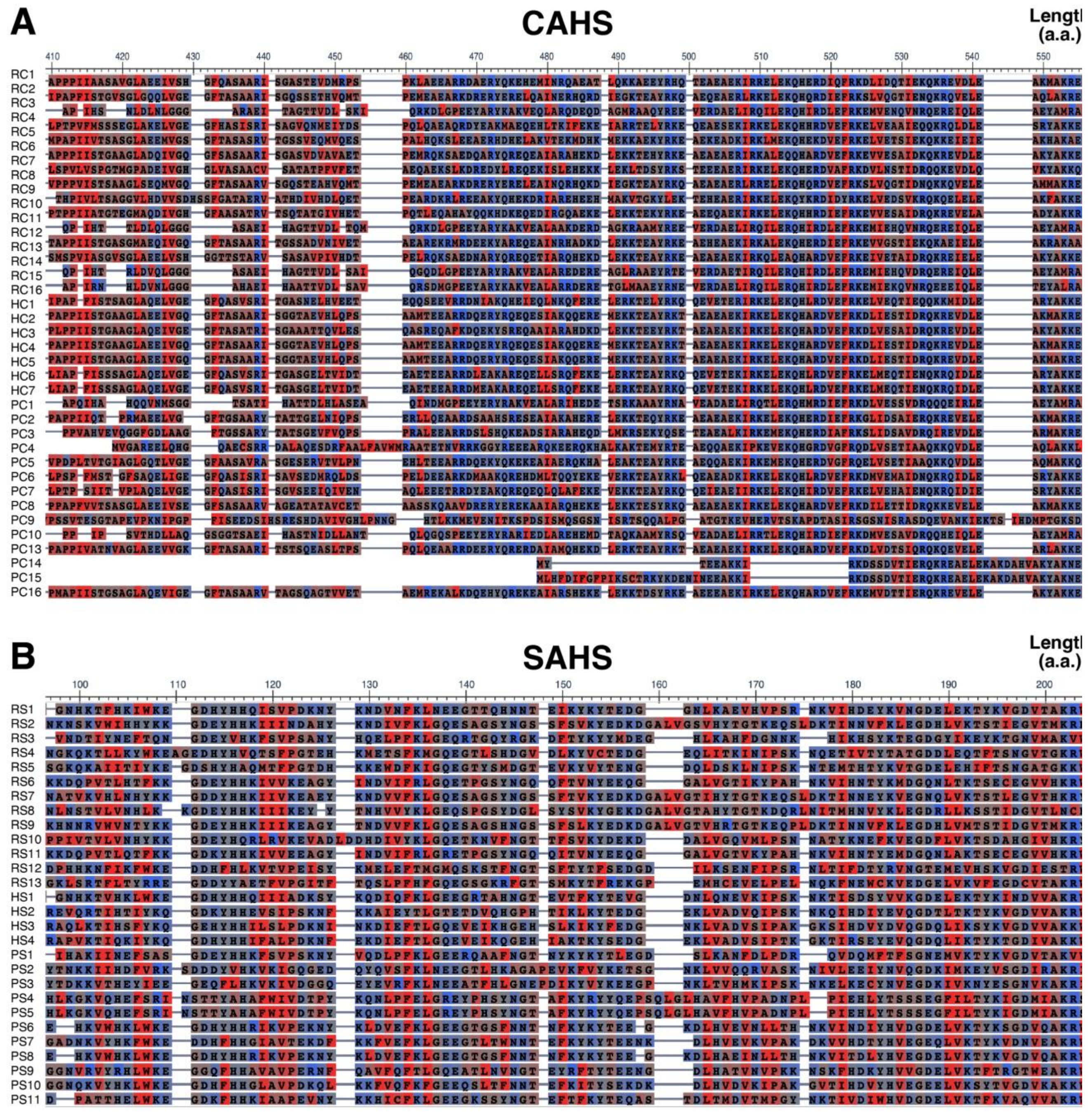


**Figure S1**. Characterization of amino acid residues within the consensus sequences of CAHSs (**A**) and SAHSs (**B**). Red, hydrophobic; blue, hydrophilic; gray, neutral.

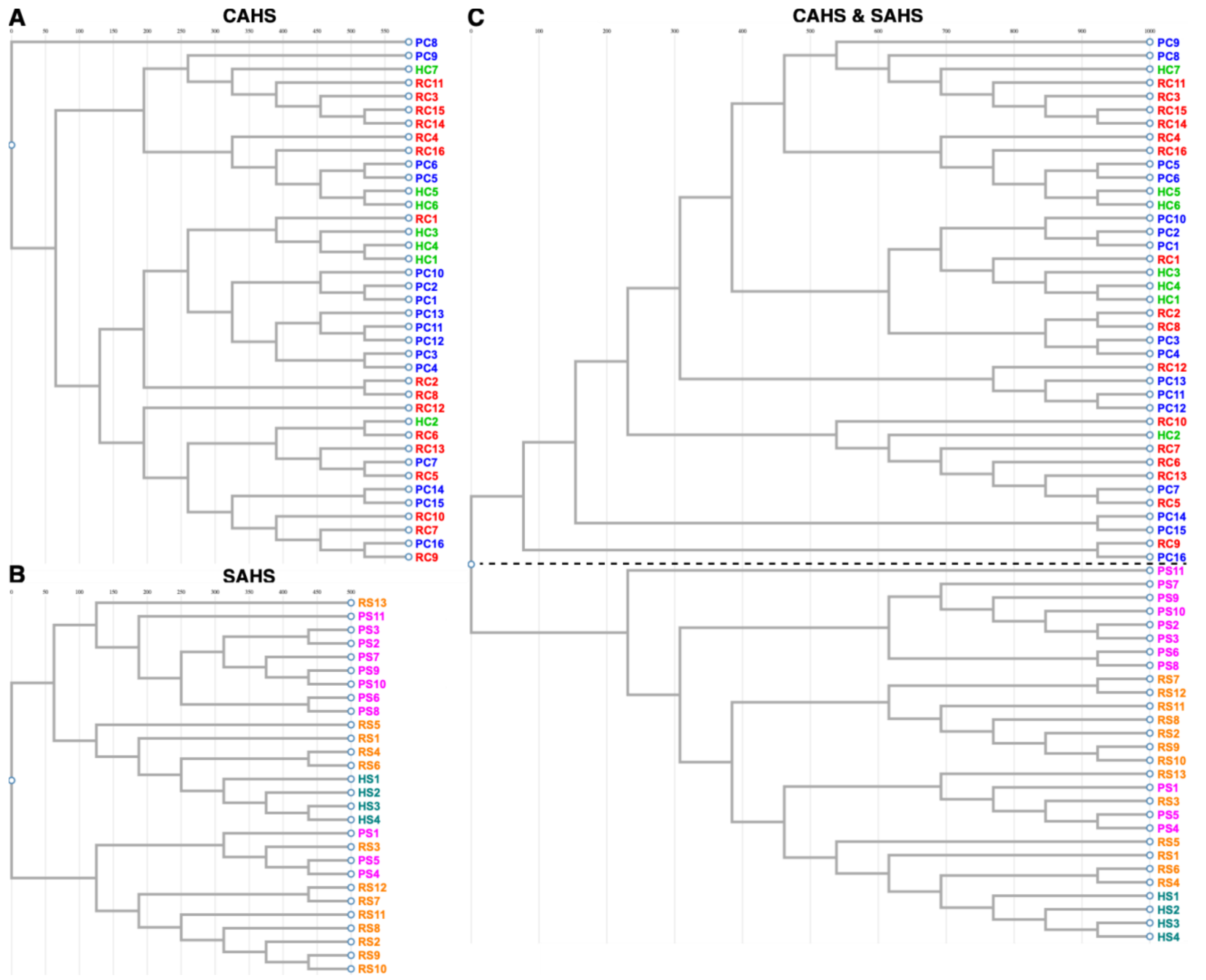


**Figure S2**. Phylogenetic trees of CAHSs (**A**), SAHSs (**B**), and both (**C**), generated by ETE3.

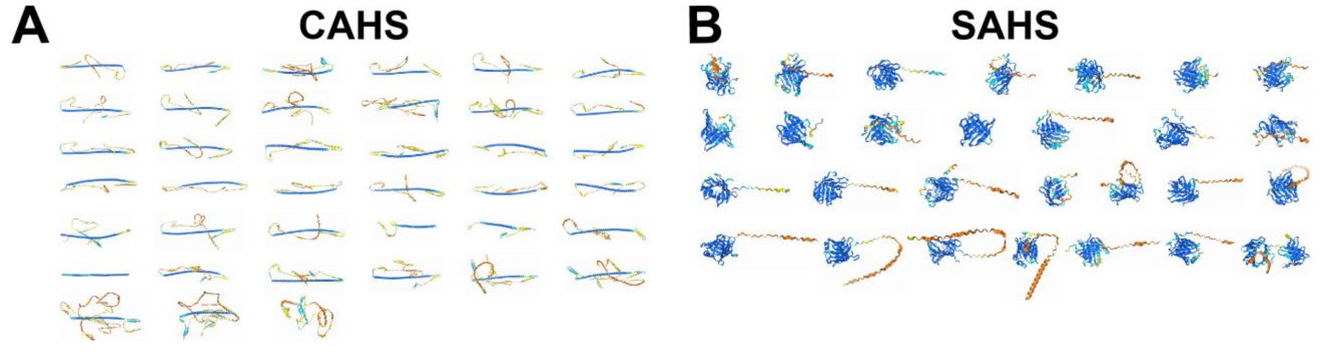


**Figure S3**. Structural prediction of 39 CAHSs and 28 SAHSs in monomer (**A** and **B**). The prediction confidence is measured as the predicted local distance difference test (plDDT). Dark blue indicates high confidence (i.e., plDDT > 90), blue to cyan indicates good confidence (i.e., 90 > plDDT > 70), yellow indicates low confidence (i.e., 70 > pTM > 50), and orange indicates very low confidence (i.e., pTM < 50).

### II. Supplementary Tables

**Table S1:** All CAHSs used in this study

| Name | GenBank ID | Abbreviation |
|---|---|---|
| **Rv CAHS1** | 402783758 | RC1 |
| **Rv CAHS2** | 402783760 | RC2 |
| **Rv CAHS3** | 402783762 | RC3 |
| **Rv CAHS4** | 1101393619 | RC4 |
| **Rv CAHS5** | 1101393709 | RC5 |
| **Rv CAHS6** | 1101393893 | RC6 |
| **Rv CAHS7** | 1101396435 | RC7 |
| **Rv CAHS8** | 1101389352 | RC8 |
| **Rv CAHS9** | 1101387114 | RC9 |
| **Rv CAHS10** | 1101380788 | RC10 |
| **Rv CAHS11** | 1101379708 | RC11 |
| **Rv CAHS12** | 1101379978 | RC12 |
| **Rv CAHS13** | 1101378429 | RC13 |
| **Rv CAHS14** | 1101375151 | RC14 |
| **Rv CAHS15** | 1101375152 | RC15 |
| **Rv CAHS16** | 1101372861 | RC16 |
| **He CAHS 94063** | 1209747827 | HC1 |
| **He CAHS 86272** | 1209747819 | HC2 |
| **He CAHS 94205 (1)** | 1209747816 | HC3 |
| **He CAHS 94205 (2)** | 1209747825 | HC4 |
| **He CAHS 77611** | 1209747824 | HC5 |
| **He CAHS 77580** | 1209747822 | HC6 |
| **He CAHS 89226** | 1209747829 | HC7 |
| **Pm CAHS 94205-like (1)** | 2493494936 | PC1 |
| **Pm CAHS 94205-like (2)** | 2493480804 | PC2 |
| **Pm CAHS 2-like** | 2493480067 | PC3 |
| **Pm CAHS 94205-like (3)** | 2493479718 | PC4 |
| **Pm CAHS 107838-like (1)** | 2493469471 | PC5 |
| **Pm CAHS 107838-like (2)** | 2493468852 | PC6 |
| **Pm CAHS 86272-like** | 2493445383 | PC7 |
| **Pm CAHS 89226-like** | 2493438951 | PC8 |
| **Pm CAHS 89226-like (2)** | 2493438943 | PC9 |
| **Pm CAHS 94205-like (4)** | 2493428999 | PC10 |
| **Pm CAHS 86272-like (X2)** | 2493423542 | PC11 |
| **Pm CAHS 86272-like (X1)** | 2493423539 | PC12 |
| **Pm CAHS 86272-like (2)** | 2493413587 | PC13 |
| **Pm CAHS 86272-like (3)** | 2493398612 | PC14 |
| **Pm CAHS 86272-like (4)** | 2493398609 | PC15 |
| **Pm CAHS 86272-like (5)** | 2493397557 | PC16 |

**Table S2:** All SAHSs used in this study

| Name | GenBank ID | Abbreviation |
|---|---|---|
| **Rv SAHS1** | 402783754 | RS1 |
| **Rv SAHS2** | 402783756 | RS2 |
| **Rv SAHS3** | 1101394461 | RS3 |
| **Rv SAHS4** | 1101395371 | RS4 |
| **Rv SAHS5** | 1101395434 | RS12 |
| **Rv SAHS6** | 1101395435 | RS5 |
| **Rv SAHS7** | 1101395438 | RS6 |
| **Rv SAHS8** | 1101395445 | RS7 |
| **Rv SAHS9** | 1101395447 | RS8 |
| **Rv SAHS10** | 1101395452 | RS9 |
| **Rv SAHS11** | 1101395453 | RS10 |
| **Rv SAHS12** | 1101395666 | RS11 |
| **Rv SAHS13** | 1101389123 | RS13 |
| **He SAHS 63681** | 1210274786 | HS1 |
| **He SAHS 53582** | 1210274784 | HS2 |
| **He SAHS 33020** | 1210274782 | HS3 |
| **He SAHS 68234** | 1210274787 | HS4 |
| **Pm SAHS 1-like (1)** | 2493482411 | PS1 |
| **Pm SAHS 1-like (2)** | 2493456970 | PS2 |
| **Pm SAHS 1-like (3)** | 2493420974 | PS3 |
| **Pm SAHS 1-like X2** | 2493398470 | PS4 |
| **Pm SAHS 1-like X1** | 2493398467 | PS5 |
| **Pm SAHS 1-like (4)** | 2493395038 | PS6 |
| **Pm SAHS 1-like (5)** | 2493392926 | PS7 |
| **Pm SAHS 1-like (6)** | 2493392853 | PS8 |
| **Pm SAHS 1-like (7)** | 2493395084 | PS9 |
| **Pm SAHS 646811-like (1)** | 2493394257 | PS10 |
| **Pm SAHS 646811-like (2)** | 2493392930 | PS11 |

**III. Supplementary Movies**

1. $Ca^{2+}$-imaging of Controls after 1, 5, and 20-minute airdry (3 AVI files)
2. $Ca^{2+}$-imaging of CAHS3-expressing cultures after 1, 5, and 20-minute airdry (3 AVI files)
3. 3D view of reconstructed confocal images showing CAHS3-AcGFP1 (green), DsRed (red), and FM4-64 (blue) in 3T3 cells after 5-minute airdry (3 AVI files)
